# Time-ResNeXt for epilepsy recognition based on EEG signals

**DOI:** 10.1101/2019.12.27.889238

**Authors:** shaoqiang Wang, Yifan Wang, Shudong Wang

**Author notes:** Corresponding author: shaoqiang Wang.

## Abstract

**Objective:** To automatically detect dynamic EEG signals to reduce the time cost of epilepsy diagnosis. In the signal recognition of electroencephalogram (EEG) of epilepsy, traditional machine learning and statistical methods require manual feature labeling engineering in order to show excellent results on a single data set. And the artificially selected features may carry a bias, and cannot guarantee the validity and expansibility in real-world data. In practical applications, deep learning methods can release people from feature engineering to a certain extent. As long as the focus is on the expansion of data quality and quantity, the algorithm model can learn automatically to get better improvements. In addition, the deep learning method can also extract many features that are difficult for humans to perceive, thereby making the algorithm more robust.

**Method:** Based on the design idea of ResNeXt deep neural network, this paper designs a Time-ResNeXt network structure suitable for time series EEG epilepsy detection to identify EEG signals.

**Results:** The accuracy rate of Time-ResNeXt in the detection of EEG epilepsy can reach 90.50%.

**Conclusion:** The Time-ResNeXt network structure produces extremely advanced performance on the benchmark dataset (Berne-Barcelona dataset), and has great potential for improving clinical practice.

## Introduction

Epilepsy is a brain disease that is caused by persistent susceptibility to recurrent seizures and the neurobiological, cognitive, psychological, and social consequences that result. According to estimates by the World Health Organization (WHO), about 2.4 million people worldwide are diagnosed with epilepsy every year [1]. Prolonged, frequent or severe seizures can lead to further brain damage and even persistent neuropsychiatric disorders. Sudden epilepsy (SUDEP) is a serious complication of epilepsy and is one of the most common causes of death in younger patients with epilepsy. The timely diagnosis of the presence and type of epilepsy is critical to its prognosis and choice of treatment options [2]. However, the diagnosis of epilepsy is relatively difficult, especially for the detection of seizures in newborns [3,4]. The usual clinical experience is judged by observing the behavior and other seizures of the newborn, but this is easily confused with other normal behaviors [5].

Epilepsy is often attributed to excessive abnormal discharges of neurons in the brain [6,7]. Electroencephalogram (EEG) signals provide a powerful tool for the diagnosis of epilepsy. Experienced neuropathologists interpret EEG signals by observing the patterns of seizures and the period of seizures, and have formulated certain international standards to find specific signal characteristics in multi-channel electroencephalography (EEG) [8,9]. Then, the condition of the patient is judged by the EEG signal rule that is manually explained. This method is relatively time-consuming and subjective, and it is objectively prone to errors [10,11]. Therefore, a suitable mechanism is needed to automatically interpret and classify EEG signals in patients with epilepsy.

EEG signal automatic classification methods usually use traditional manual feature machine learning and statistical methods, such as time-frequency analysis using wavelet transform [12], detection method using entropy estimator [13], discrete wavelet transform and approximate entropy Method [14] and so on. In addition, there are also methods for detecting using shallow neural networks using artificial features, such as Elman and Probabilistic Neural Networks [15], which use approximate entropy as input features of the network, artificial neural networks [16] The method using Volterra system and cellular nonlinear network [17] and so on.

With the development of the field of machine learning in recent years, a large number of excellent machine learning classification algorithms have emerged, the most representative of which is the deep neural network algorithm. Especially in the field of image classification, deep learning methods, such as VGG [18] network, Google Inception [19] network, and ResNet [20] network, have powerful automatic feature extraction capabilities [21], which have been completely completed in some fields Beyond traditional machine learning and statistical methods and shallow artificial neural network methods, it can even identify targets that are difficult to distinguish with the naked eye, surpassing humans. In addition, many large companies have also adopted the method of deep learning as one of their core competitiveness [22,23,24].

This paper draws on the excellent deep neural network structure in the image field, and designs an excellent end-to-end network structure based on ResNeXt [25] and suitable for EEG signal epileptic detection. And the performance of the network is verified on a public standard dataset (Berne Barcelona EEG dataset [26]), and compared with traditional algorithms [27,28,29,30] using this dataset, for us the performance of the algorithm is evaluated.

## 1 data preparation

### 1.1 data description

The data are from the EEG database of Berne Barcelona and are divided into two categories: EEG signal data during the onset of epilepsy patients and EEG signal data during the onset of epilepsy patients. Each category has 3750 pieces of data, each piece of data has 2 signal channels with a length of 10240 and a sampling frequency of 512Hz (the time length of each piece of data is 20s). Part of the original EEG image is shown in Figure 1.

**Figure 1.**
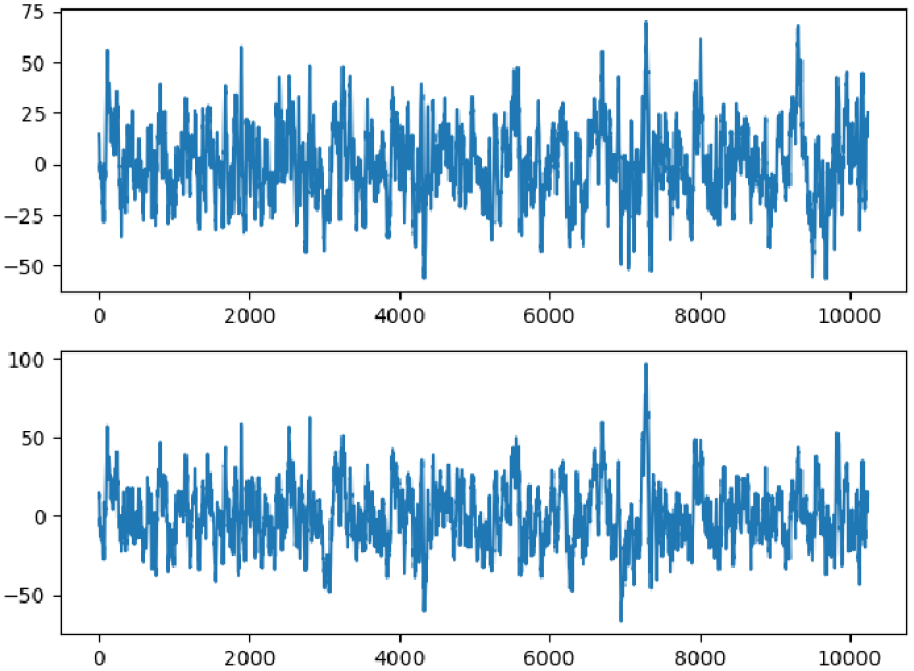
Part of the EEG image.

### 1.2 Training data preparation

Due to the balanced data classification, there is no data deviation. So there is no data enhancement for a single category. Only the data set is divided to prepare for model training. The method is shown in Table 1. The original data set is randomly divided into a training set (3000 items / category), a validation set (250 items / category), and a test set (500 items/category).

**Table 1.** Data allocation table.

|  | training set | validation set | test set |
| --- | --- | --- | --- |
| Epilepsy signal | 3000 | 250 | 500 |
| Non-epileptic signal | 3000 | 250 | 500 |
| total | 6000 | 500 | 1000 |

## 2 network model design

### 2.1 Model design ideas

ResNeXt’s deep learning network model structure design idea is followed. According to the data characteristics of EEG signals, a network structure Time-ResNeXt is designed for EEG time series classification.

According to the traditional idea of designing network structures to improve the accuracy of the model, most of them are to deepen or widen the structure of the network, but as the number of hyperparameters (such as the number of channels, the size of the convolution kernel, etc.) increases, neural network design The difficulty and computational overhead also increase greatly. The algorithm in this paper benefits from the repeated topology of the ResNeXt network sub-modules, which enables it to have a very high accuracy rate while slightly increasing the amount of network calculations, while also greatly reducing the number of hyperparameters.

First, I have to mention the classic VGG network and Inception network. The design idea of VGG network is: modularize the neural network to increase the depth, but such a deep network will cause network degradation due to gradients. The structure of VGG network key modules is shown in Figure 2.

**Figure 2.**
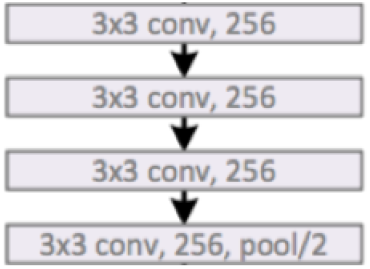
Structure of key modules of VGG network.

The design philosophy of the Inception network is exactly the opposite: the width of the network is increased by the split-transform-merge method, but the settings of the various hyperparameters of this Inception network are more targeted and need to be performed when applied to other data sets. Many modifications, so scalability is average. The structure of the key modules of the Inception network is shown in Figure 3.

**Figure 3.**
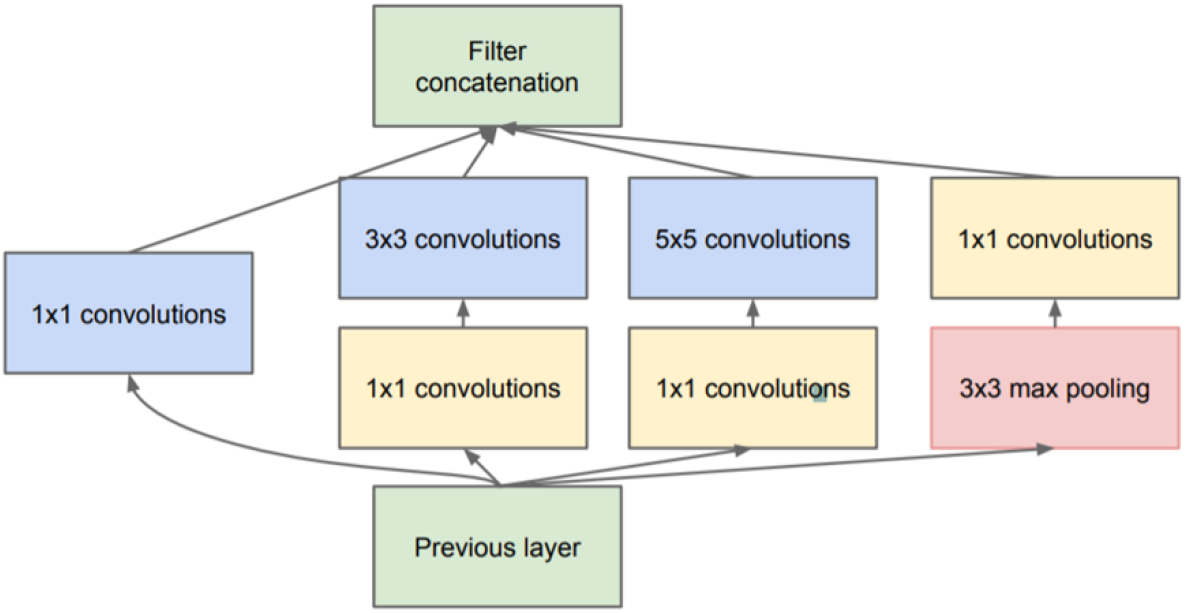
The structure of the key modules of the Inception network.

The ResNeXt network is based on the design idea of ResNet’s cross-layer connection, and combines the VGG and Inception networks. And through the structure of ResNet cross-layer connection to improve the shortcomings of VGG network too deep degradation. The cross-layer connection structure is shown in Figure 4.

**Figure 4.**
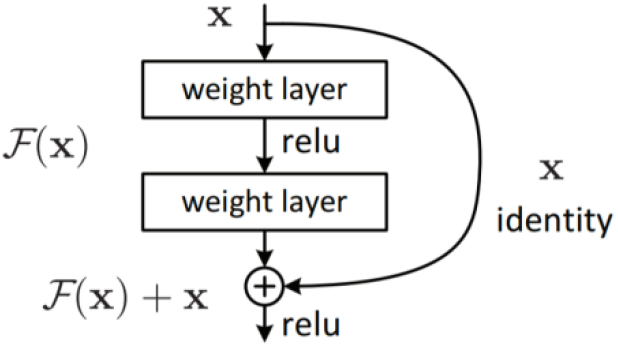
Cross-layer connection structure.

The transformation set structure is shown in Figure 5.

**Figure 5.**
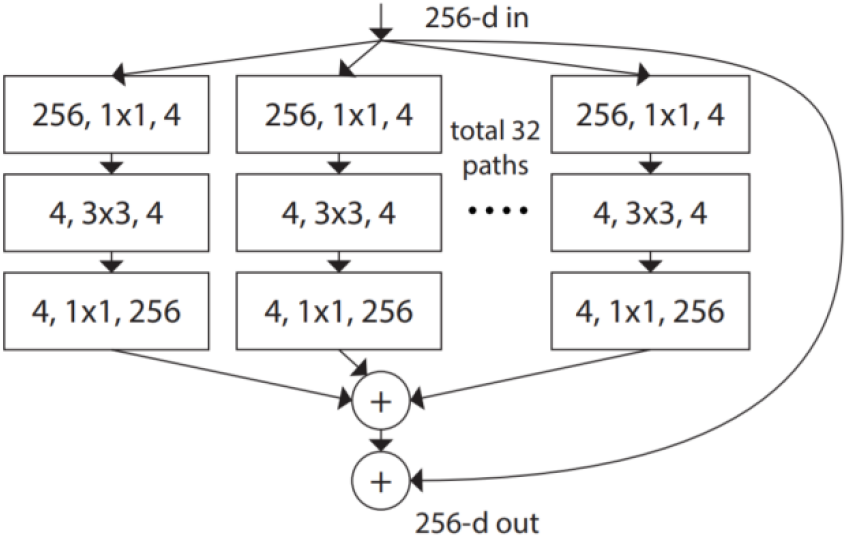
Transform Set Structure.

The convolution modules of the transform set are all the same. ResNeXt uses a transformation set to replace the transformation structure of the Inception network. Because each aggregated topology is the same, the network no longer needs to modify too many hyperparameters on different data sets, which has better robustness.

### 2.2 Model design process

1. The original ResNeXt-50 has five stages and a large number of parameters, as shown in Figure 6.
2. During training, it is found that the results are difficult to converge and tend to be completely random. Therefore, it was determined that the network structure was too complicated. Starting from the complexity of the network, the network was tailored to try to find a suitable structure. The test results are shown in Table 2.

**Figure 6.**
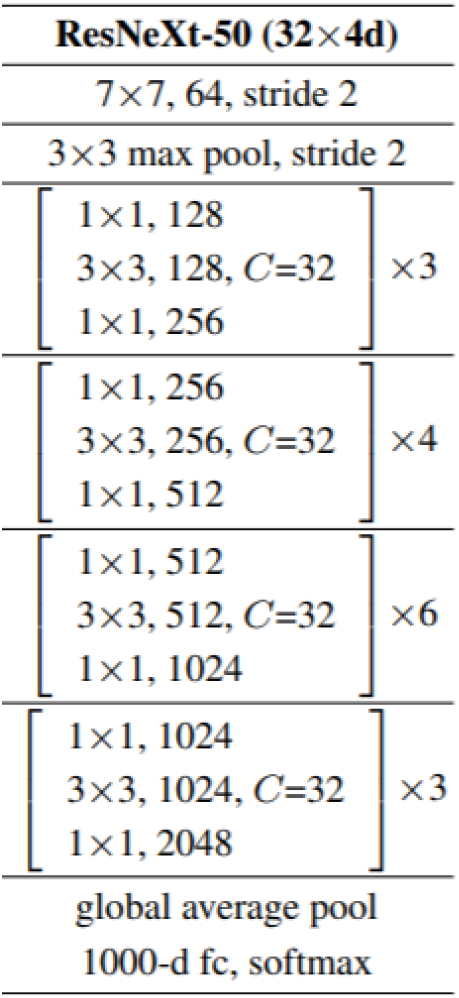
ResNeXt-50 module.

**Table 2.**
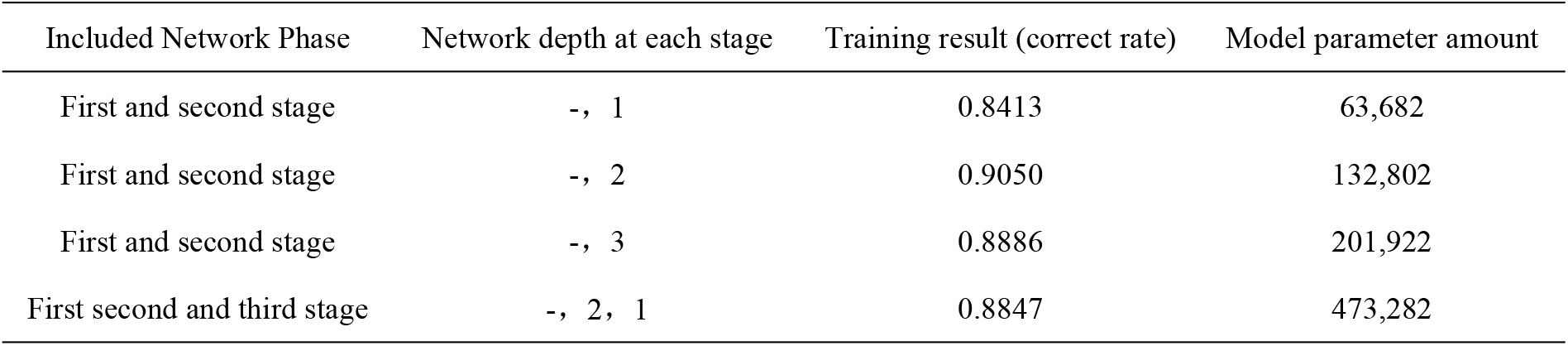
Model optimization table.

| Included Network Phase | Network depth at each stage | Training result (correct rate) | Model parameter amount |
| --- | --- | --- | --- |
| First and second stage | -, 1 | 0.8413 | 63,682 |
| First and second stage | -, 2 | 0.9050 | 132,802 |
| First and second stage | -, 3 | 0.8886 | 201,922 |
| First second and third stage | -, 2, 1 | 0.8847 | 473,282 |

Through the above experiments, the layers and depth of the network are continuously explored to train the model. Finally, it is concluded that a ResNeXt network with two stages and a depth of 2 in the second stage has the best performance, namely the final structure of Time-ResNeXt.

### 2.3 Time-ResNeXt network structure

The structure of Time-ResNeXt neural network is shown in Figure 7.

**Figure 7.**
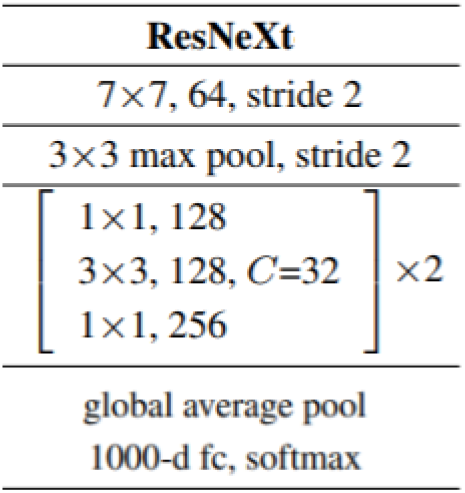
Time-ResNeXt neural network structure.

It has two phases in total. The detailed network structure of the first phase is shown in Figure 8.

**Figure 8.**
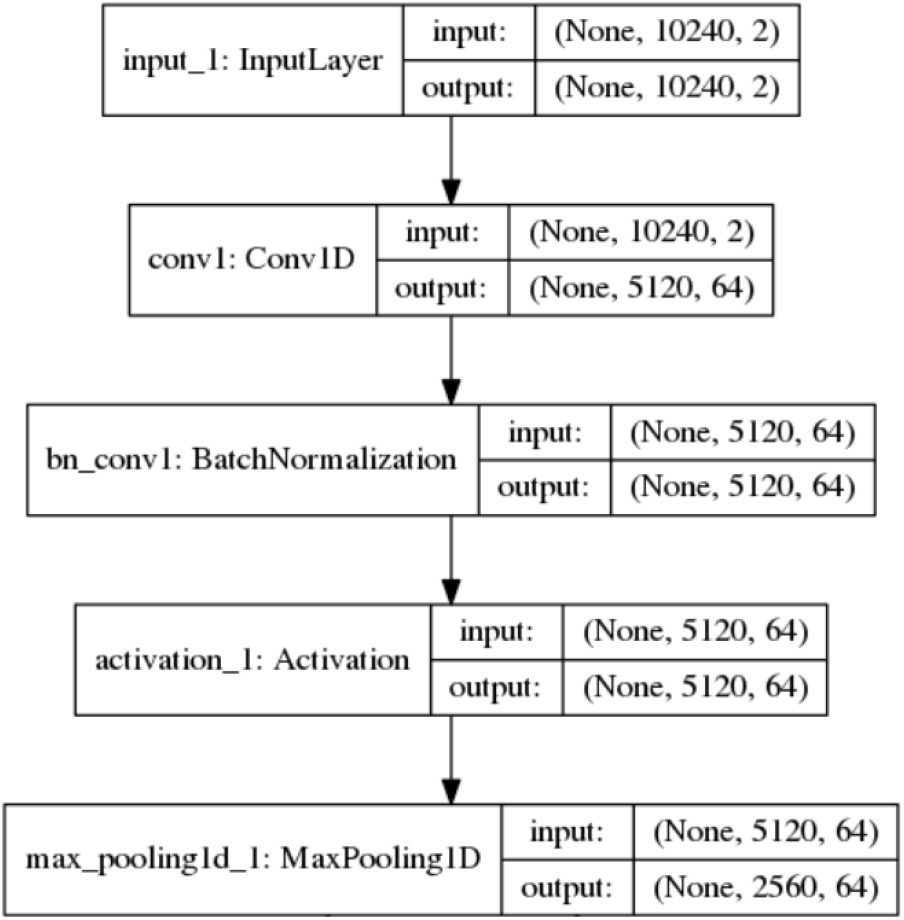
Detailed Network Structure of Time-ResNeXt Phase I.

The depth of the second phase of the network structure is 2, that is, two network structure sub-modules, each of which contains cross-layer connections, activation layers, convolutional layers, batch normalization layers, and transform set modules. The main structure is The transformation set module, which uses a network design structure in a network, is a module for forming a convolution transformation set by connecting 32 convolutional structural blocks as shown in FIG. 9 in parallel, which is the main feature extraction module.

**Figure 9.**
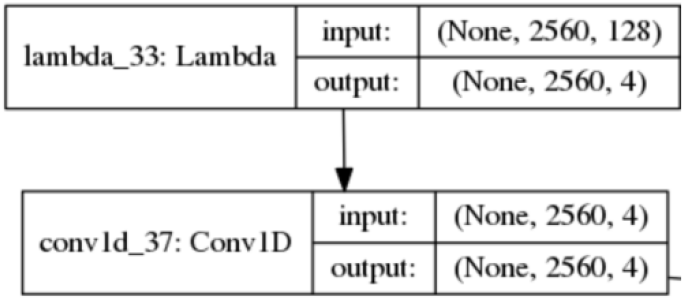
Time-ResNeXt detailed network structure in the second phase.

## 3 model training

### 3.1 Optimizer

Use Adam’s algorithm as the optimizer. The Adam algorithm is an algorithm that performs a stepwise optimization on a random objective function. This algorithm is based on adaptive low-order moment estimation, has high computational efficiency and low memory requirements. The adaptive learning rate of different parameters can be calculated by estimating the first and second gradients. In addition, the gradient rescaling of Adam’s algorithm is invariant, so it is very suitable for solving problems with large-scale data or parameters.

The advantages are: easy to implement, efficient calculation, less memory required, invariance of gradient diagonal scaling, and only minimal tuning.

The parameter settings of the Adam optimizer are shown in Table 3.

**Table 3.** Adam optimizer parameters.

| parameter name | Parameter value |
| --- | --- |
| Lr | 0.001 |
| beta_1 | 0.9 |
| beta_2 | 0.999 |
| Epsilon | 1e-8 |
| Decay | 0.01 |

Among them, lr refers to the step size, that is, the step size of each gradient descent. Decay is a weight decay factor, which avoids overfitting by adding a regular term to the loss function.

### 3.2 Loss function and evaluation index

The loss function uses the cross entropy function of L2 regularization attenuation. The cross entropy function is calculated as follows.

Among them, is the true label of each instance, and is the predicted probability value of each instance. Then, add regularized attenuation to the loss function to avoid overfitting. The method is shown in the following formula.

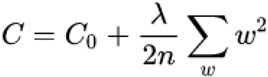

The first term *C*_0_ is the original loss function, which is the cross-entropy function. The second term λ is a regular term, a regular term coefficient, n is the number of training samples, and w is a parameter of the network. Weight decay (L2 regularization) can effectively prevent overfitting.

The evaluation index adopts the correct rate index. Here, the concepts of TP (true), TN (true negative), FP (false positive), and FN (false negative) are introduced first. The classification of evaluation indicators is shown in Figure 10.

**Figure 10.**
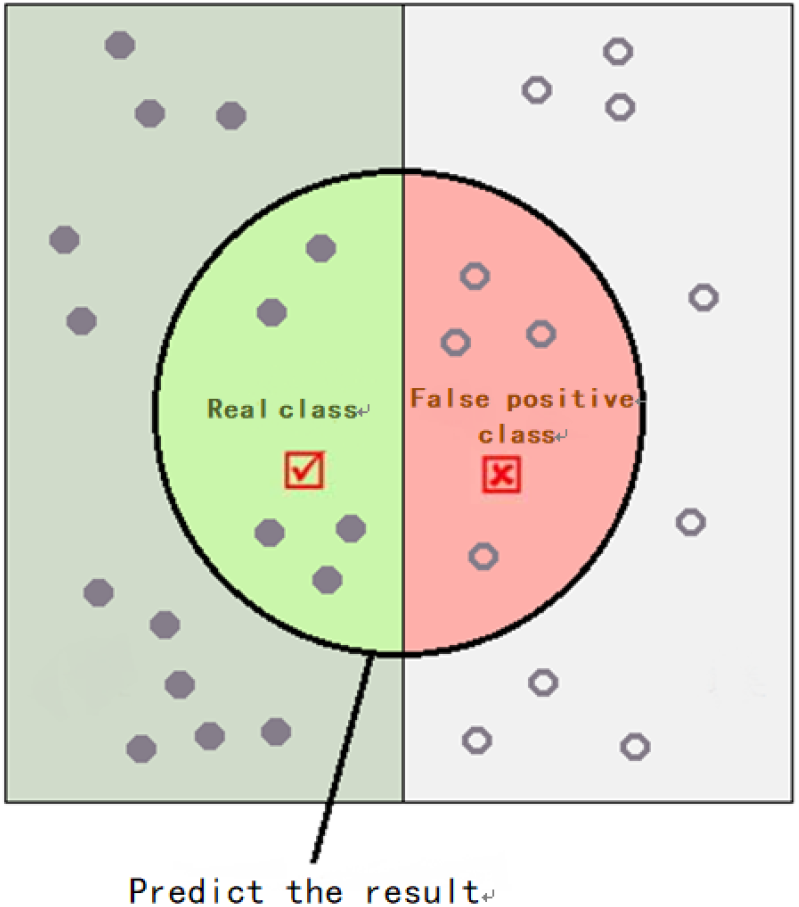
Classification of evaluation indicators.

the accuracy calculation method is:

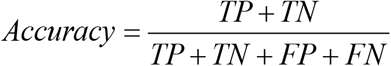

### 3.3 Early stop

In order to further avoid overfitting, an early stopping training strategy is adopted. When the model exceeds 30 consecutive generations of evaluation indicators and does not improve on the validation set, training is stopped. This can prevent the model from over-learning on the training set, avoid excessive bias, and reduce the generalization performance of the model.

### 3.4 Training process records

The results of the training set are shown in Figure 11, (the X-axis is the training algebra, and the Y-axis is the training evaluation index)

**Figure 11.**
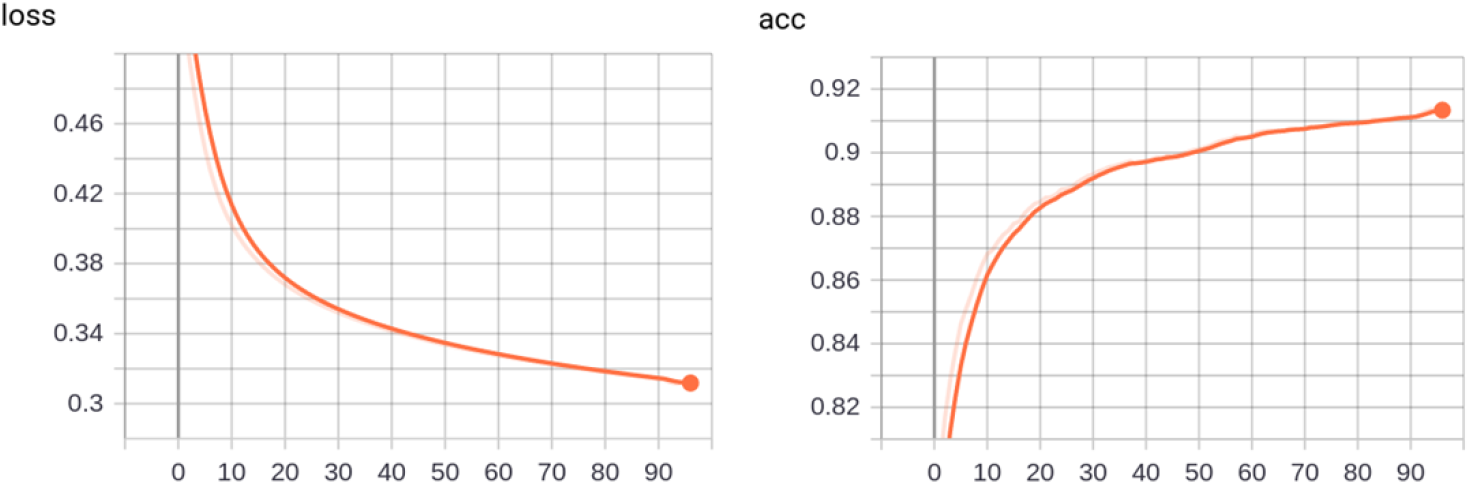
Training set results.

The results of the validation set are shown in Figure 12.

**Figure 12.**
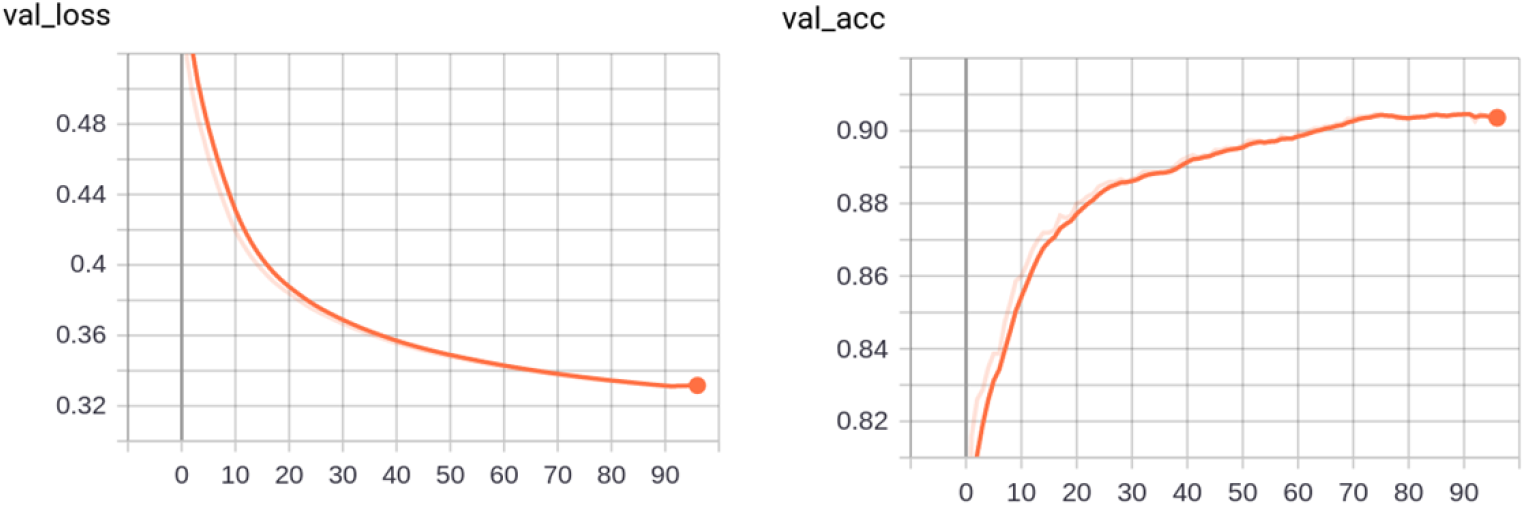
validation set results.

The results show that at 74th generation, the model performs best on the validation data, with a correct rate of 0.9050.

## 4 results and discussion

Through continuous training of the model, the accuracy rate finally reached 0.9050, achieving an extremely advanced performance. Its pair is shown in Table 4.

**Table 4.**
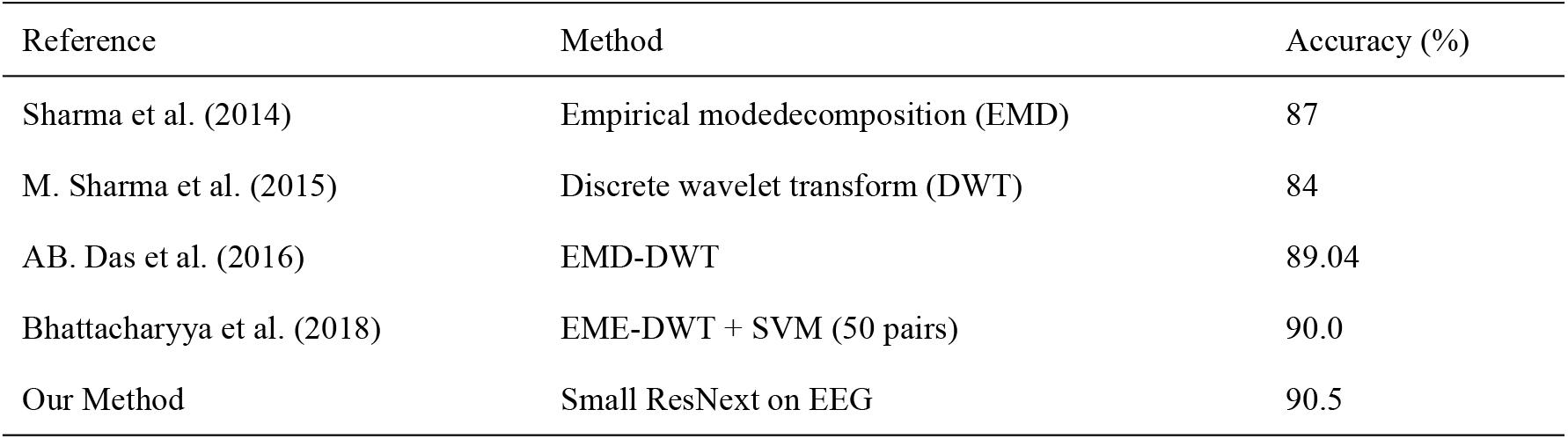
Model comparison.

| Reference | Method | Accuracy (%) |
| --- | --- | --- |
| Sharma et al. (2014) | Empirical modedecomposition (EMD) | 87 |
| M. Sharma et al. (2015) | Discrete wavelet transform (DWT) | 84 |
| AB. Das et al. (2016) | EMD-DWT | 89.04 |
| Bhattacharyya et al. (2018) | EME-DWT + SVM (50 pairs) | 90.0 |
| Our Method | Small ResNext on EEG | 90.5 |

Other related evaluation indicators are shown in Table 5.

**Table 5.**
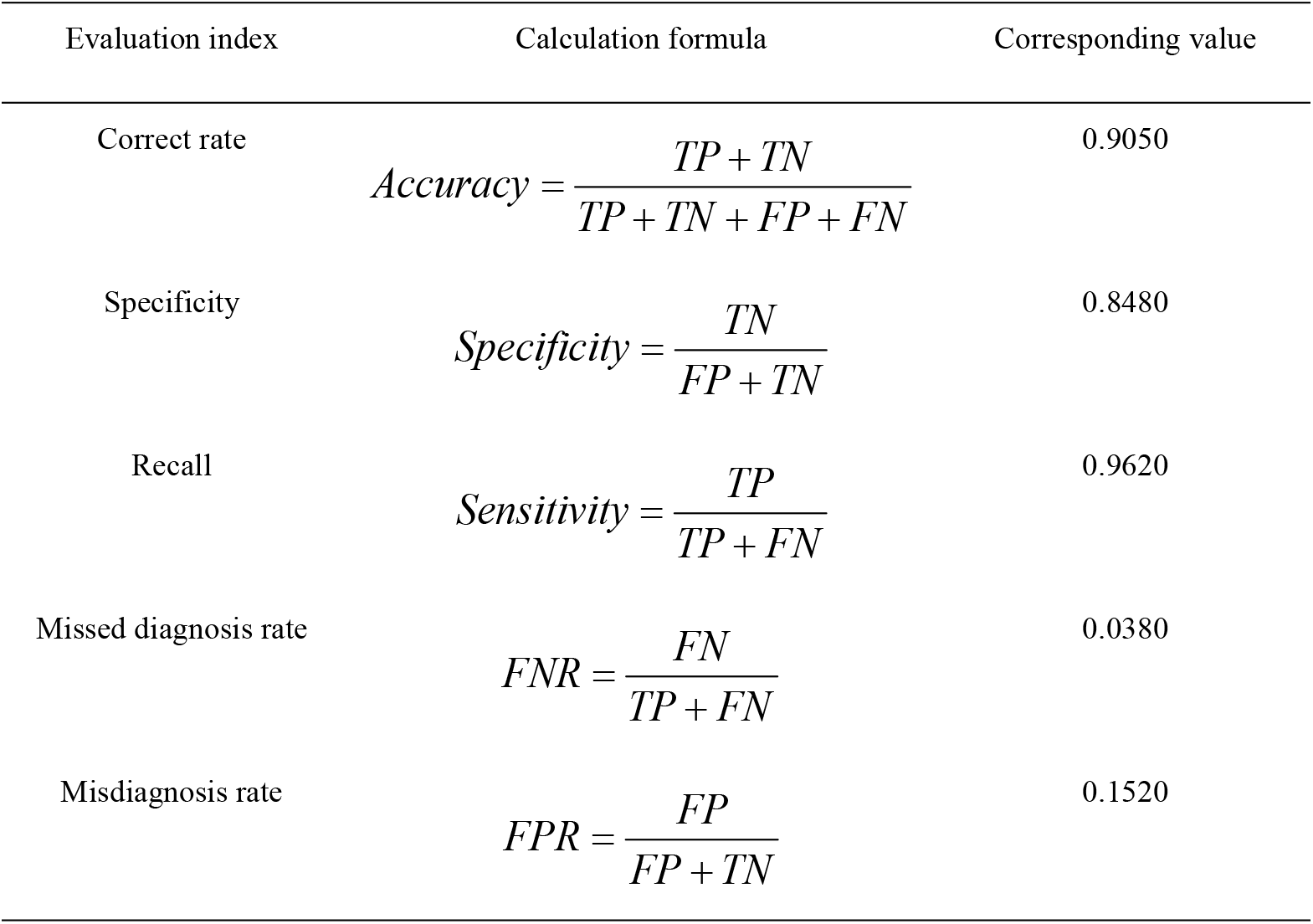
Time-ResNeXt network evaluation.

In addition, there are many areas in this mission where you can continue to improve. For example, you can use Hard Example Mining to train difficult samples. Increasing the amount of data is also an extremely good method. The causes of epilepsy are complex. Trying more detailed multi-classification will help decouple the data and further reduce the difficulty of model training.

All in all, our classifier has achieved an extremely good performance and has excellent scalability on the Bern dataset in Barcelona. As the amount of data in real business scenarios increases, it will show even better performance.

## Compliance with Ethical Standards

`Ethical approval: This article does not contain any studies with human participants or animals performed by any of the authors`.

`Disclosure of potential conflicts of interest: NONE`

`This work was supported by the National Natural Science Foundation of China` (61873281, 61572522, 61502535, 61972416 `and` 61672248).

